# Re-assessment of the subcellular localization of Bazooka/Par-3 in *Drosophila*: No evidence for localization to the nucleus and the neuromuscular junction

**DOI:** 10.1101/2023.12.21.572772

**Authors:** Soya Kim, Jaffer Shahab, Andreas Wodarz

## Abstract

Bazooka/Par-3 (Baz) is an evolutionarily conserved scaffold protein that together with its binding partners atypical protein kinase C (aPKC) and Par-6 forms the Par complex. This protein complex functions as a master regulator for the establishment and maintenance of cell polarity in many different cell types. In the vast majority of published research papers Par complex members have been reported to localize at the cell cortex and at intercellular junctions. However, there have also been several reports claiming localization and function of Par complex members at additional subcellular sites, in particular the nucleus, the nuclear envelope and the neuromuscular junction. In this study we have re-assessed this issue for Baz in a systematic manner. We used a variety of antibodies raised in different host animals against different epitopes of Baz for confocal imaging of *Drosophila* tissues. We tested the specificity of these antisera by using mosaic analysis with null mutant *baz* alleles and tissue-specific RNAi against *baz.* In addition, we used a GFP-tagged gene trap line for Baz and a bacterial artificial chromosome (BAC) expressing GFP-tagged Baz under control of its endogenous promoter in a *baz* null mutant background to compare the subcellular localization of the respective GFP-Baz fusion proteins to the staining results with anti-Baz antisera. Together, these experiments did not provide any evidence for specific localization of Baz to the nucleus or the neuromuscular junction.

## Introduction

Baz/Par-3 is a multi-domain scaffold protein with well characterized roles in the establishment and maintenance of cell polarity. Baz is a core member of the polarity regulating Par complex, which it forms together with its evolutionarily conserved binding partners aPKC and Par-6 (Goldstein and Macara, 2007; Johnson and Wodarz, 2003; Johnston and Ahringer, 2010; Tepass, 2012). Baz/Par-3 was first identified in *C. elegans* (Etemad-Moghadam et al., 1995) with the subsequent identification of homologues in *Drosophila* (Kuchinke et al., 1998), and mammals (Izumi et al., 1998). In *C. elegans*, Baz/Par-3 functions as a key regulator of spindle orientation and polarity in the early embryo (Etemad-Moghadam et al., 1995) and the assembly of adherens junctions in the intestine (Achilleos et al., 2010). In *Drosophila*, Baz has been shown to function as a key regulator of apico-basal polarity in epithelial cells (Harris, 2012; Tepass, 2012). Baz functions at the top of a genetic hierarchy in the establishment of adherens junctions during cellularization (Harris and Peifer, 2004; Harris and Peifer, 2005) and in the maintenance of apico-basal polarity during embryonic epithelial development (Bilder et al., 2003; Müller and Wieschaus, 1996; Tanentzapf and Tepass, 2003). Baz is also required for oocyte differentiation (Cox et al., 2001) and polarization of the developing oocyte along the anterior/posterior axis (Doerflinger et al., 2010). In the developing nervous system, Baz is essential for the maintenance of stem cell fate in neuroblasts through the establishment of apico-basal polarity, spindle orientation and asymmetric cell division (Kuchinke et al., 1998; Loyer and Januschke, 2020; Schober et al., 1999; Wodarz et al., 1999). In mammalian epithelial cells, Par-3 has been shown to play a crucial role in the assembly and maintenance of tight junctions and adherens junctions as well as epithelial spindle orientation (Chen and Macara, 2005; Hao et al., 2010; Ooshio et al., 2007).

Aside from their cortical localization and roles in regulating cell polarity, several studies have reported that members of the Par complex also localize to the nucleus (Cline and Nelson, 2007; Fang et al., 2007; Perander et al., 2000; Seidl et al., 2012; Speese et al., 2012). Studies in mammalian cell culture have revealed that in response to DNA damage induced by γ-irradiation, Par3 translocates to the nucleus where it associates with DNA-dependent protein kinase (DNA-PK) to mediate DNA double strand break repair (Fang et al., 2007). A study by Speese et al., 2012 revealed that Par proteins also show nuclear localization in *Drosophila* body wall muscles where they promote neuro-muscular junction (NMJ) formation. They reported that Baz localizes to the nuclear envelope, including nuclear envelope foci containing a C-terminal cleavage product of the *Drosophila* Wingless/Wnt1 receptor DFrizzled 2 (DFz2C). At these foci, Baz is required for phosphorylation of the nuclear envelope component LaminC (LamC) by aPKC to promote nuclear envelope budding, facilitating the export of ribonucleoprotein particles containing DFz2C and mRNAs encoding post-synaptic proteins (Speese et al., 2012).

In this study, we sought to further analyze Baz nuclear envelope localization in *Drosophila* to gain insight into any additional functions it may have in the nucleus. By immunostaining analysis and two independent Baz-GFP fusion protein lines (Besson et al., 2015; Buszczak et al., 2007) under the endogenous promotor, we assessed whether Baz nuclear envelope localization was ubiquitous or restricted to specific cell types or developmental stages. Immunostaining analysis using multiple antibodies raised against different domains of Baz revealed that immunostaining for Baz at the nuclear envelope was detectable in some but not all tissues, in particular in polyploid cells. However, to our surprise mutational analysis revealed that Baz nuclear envelope immunostaining persisted in *baz* mutant clones although cortical staining was completely abandoned. Furthermore, analysis of Baz-GFP fusion protein lines showed no Baz-GFP nuclear envelope localization. The same was true for localization of Baz to the NMJ. Thus, our results provide strong evidence that Baz does neither localize to the nuclear envelope nor to the NMJ, requiring a reassessment of its reported function at these subcellular sites.

## Results

### Baz localization at the nuclear envelope is observed with different anti-Baz antibodies

We assessed by indirect immunofluorescence whether Baz also localizes to the nucleus, in addition to its well-described localization to the cytocortex and intercellular junctions. Several antibodies were used to stain fat body tissue dissected from wild type 3^rd^ instar larvae. Antibodies raised in rabbit and rat against the Baz N-terminal region (amino acids 1-297; (Wodarz et al., 1999; Wodarz et al., 2000)), an antibody raised in guinea pig against the Baz PDZ domains (amino acids 309-747; (Shahab et al., 2015)) and an antibody raised in guinea pig against a region in the C-terminal half of Baz (amino acids 905-1221, this work) all showed nuclear staining (Fig. S1). For all further analyses we made use of the antibody against the N-terminal region of Baz raised in rabbits, as in the *Drosophila* community it is the most commonly used antibody to detect Baz. Also, it showed the least background staining in comparison to the other antibodies tested. Closer inspection of the immunofluorescence pattern detected by this antibody in fat body showed a localization to the periphery of the nucleus colocalizing with the nuclear envelope component Lamin C (Fig. 1A), indicating that Baz localizes to the nuclear envelope.

**Figure 1.**
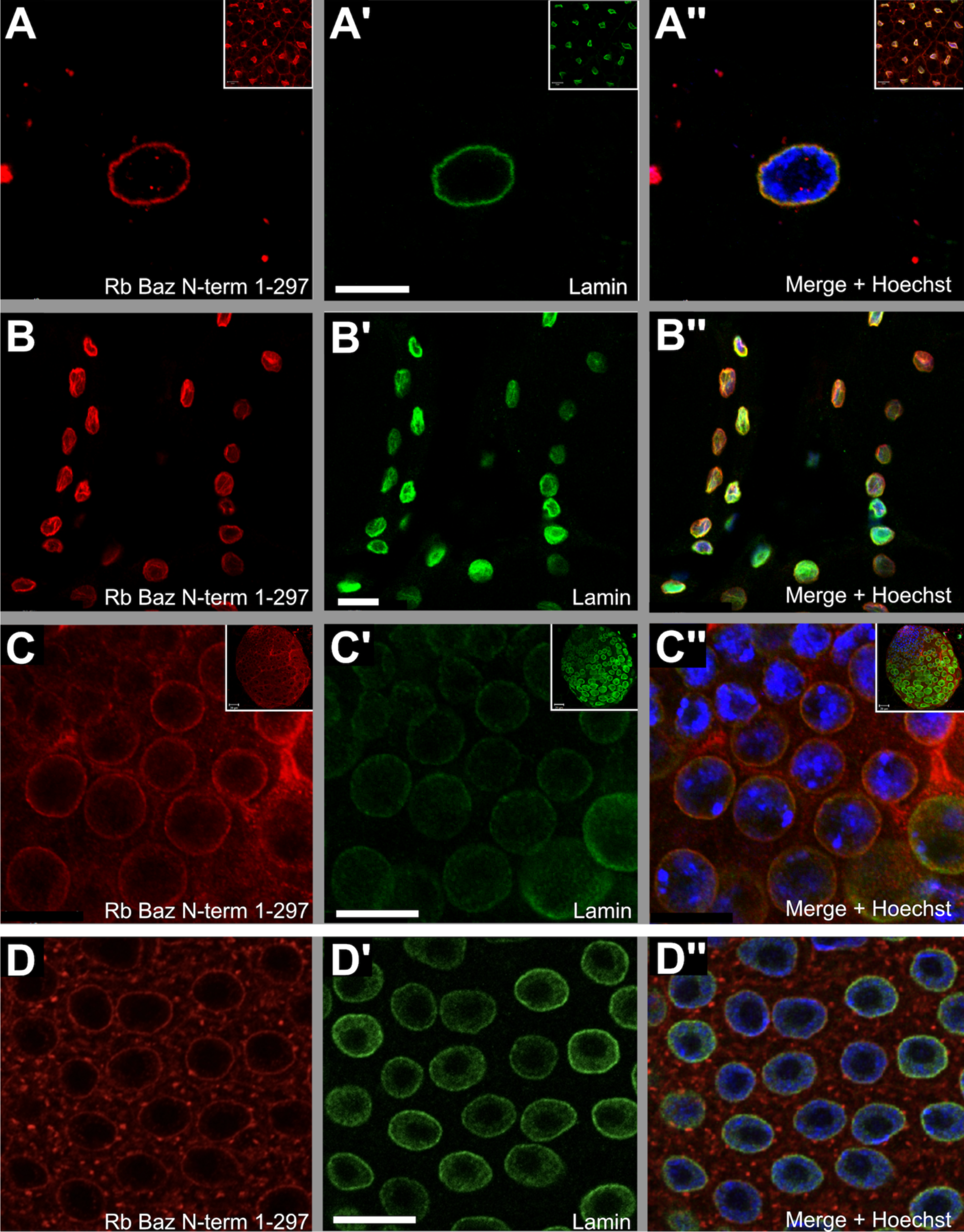
Bazooka immunostaining colocalizes with Lamin C at the nuclear envelope of 3^rd^ instar fat body, body wall muscle, spermatogonia and adult follicular epithelial cells. (A-A’’) Fat body tissue, (B-B’’) body wall muscles and (C-C’’) testis dissected from 3^rd^ instar wild type larvae were stained with rabbit anti-Bazooka 1-297 (anti Baz, red), Lamin C (green) to mark the nuclear envelope and Hoechst (blue) to label DNA. The high magnification image of anti-Baz immunostaining in adipocytes (A) shows a strong signal in the periphery of the nucleus in a pattern identical to Lamin C (A’). The merged image shows colocalization of both signals (A’’). Insets in (A-A’’) show low magnification images of the immunostained fat body tissue. Body wall muscles show a similar pattern of nuclear envelope localization for anti-Baz immunostaining (B), Lamin C (B’) and their corresponding colocalization (B’’). (C-C’’) Nuclear envelope localization for anti-Baz immunostaining is also observed in clusters of spermatogonia in larval testis (C) where it colocalizes with Lamin C (C’, C’’). Insets in (C-C’’) show low magnification images of larval testis. (D-D’’) Adult follicular epithelial cells stained with anti-Baz (D) also show a weak signal at the nuclear envelope that colocalizes with Lamin C (D’, D’’). Scale bars in A-A’’ and C-D’’= 10 µm, in B-B’’ = 20 µm.

### Nuclear envelope immunofluorescence for Baz is observed in different tissues

We next sought to determine whether Baz nuclear envelope localization occurred in a tissue-specific manner. In addition to larval fat body cells (Fig. 1A-A’’, Fig. S1), we observed Baz nuclear envelope localization in larval body wall muscle (Fig. 1B-B’’), spermatogonial clusters in larval testis (Fig. 1C-C’’), adult ovary follicular epithelial cells (Fig. 1D-D’’), nurse cells (Fig S2A-A’’) and oocytes (Fig. S2B-B’’). Baz nuclear envelope immunostaining signal was absent in larval wing imaginal disc cells (Fig. S2C-C’’) and in embryonic tissue, e. g. epidermis (Fig. S2D-D’’) and neuroblasts (Fig. S2D-D’’).

### Fully functional Baz-GFP fusion proteins do not show nuclear envelope localization

While strong nuclear envelope immunostaining was observed using several independently raised anti Baz antibodies (Fig. 1; Fig. S1; Fig. S2; Fig. S3), no nuclear localization was detected using a Baz-GFP BAC line (Besson et al., 2015) (Fig. S3C-D’’) nor in a GFP-Baz protein-trap line (Buszczak et al., 2007). In the protein-trap line an engineered exon encoding for GFP is spliced in between the first and third exon (the second exon is excluded in all isoforms). As two transcripts start their translation within the third exon, it is likely that the other two transcripts starting in exon 1 translate the GFP to generate the fusion protein. However, flies expressing this GFP-Baz under the endogenous promotor are homozygous viable and are phenotypically indistinguishable from wild type flies, indicating that this construct is fully functional. The BAC line integrates the GFP within exon 10, giving rise to GFP fusion proteins for all four isoforms. Like in the protein-trap line, the BAC line in the *baz^EH747^* null mutant background is homozygous viable and therefore the construct fully functional (Fig. S4D-D’’). For both lines Baz-GFP expression was detected at the junctions, colocalizing with the signal from the Baz N-terminal antibody (Fig. S3C, C’, E, E’, Fig. S4D-E’’). However, even after signal enhancement using an anti-GFP antibody we did not detect a signal at the nuclear envelope in follicular epithelial cells like seen when using the anti-Baz antibodies (Fig. S3D, D’, F, F’), nor at the nuclear envelope of the oocyte (Fig. S4D-E’’).

### Baz nuclear envelope immunostaining in the ovary is an artifact

In the follicular epithelium, Baz immunostaining detected Baz at the adherens junctions (Fig. S3A) and in a different focal plane at the nuclear envelope (Fig. S3B). To test whether both junctional and nuclear envelope staining of Baz was specific, we eliminated *baz* expression in the follicular epithelium using two different methods. First, we used RNAi against *baz* driven by *traffic jam* (*tj*)::Gal4, which is expressed in all follicle cells of the developing egg chamber until stage 12 (Fig. 2A’) (Li et al., 2003). During early stages of egg chamber development the expression of *tj::*Gal4 in the follicle cells is heterogeneous (Fig. 2A’), while it becomes homogeneous in older egg chambers (inset in Fig. 2A’). Whereas the junctional staining for Baz close to the apical surface of the follicular epithelium was lost upon RNAi against *baz* (Fig. 2B), the staining at the nuclear envelope persisted in the same egg chamber imaged at the optical plane containing the nuclei (Fig. 2C). This result was confirmed using the CY2::Gal4 driver line expressed in the follicular epithelium and with three different RNAi lines against *baz* (data not shown). The staining for Armadillo/beta-catenin (Arm) (Fig. 2B’, C’) shows junctions that are still intact although Baz is downregulated (Shahab et al., 2015).

**Figure 2.**
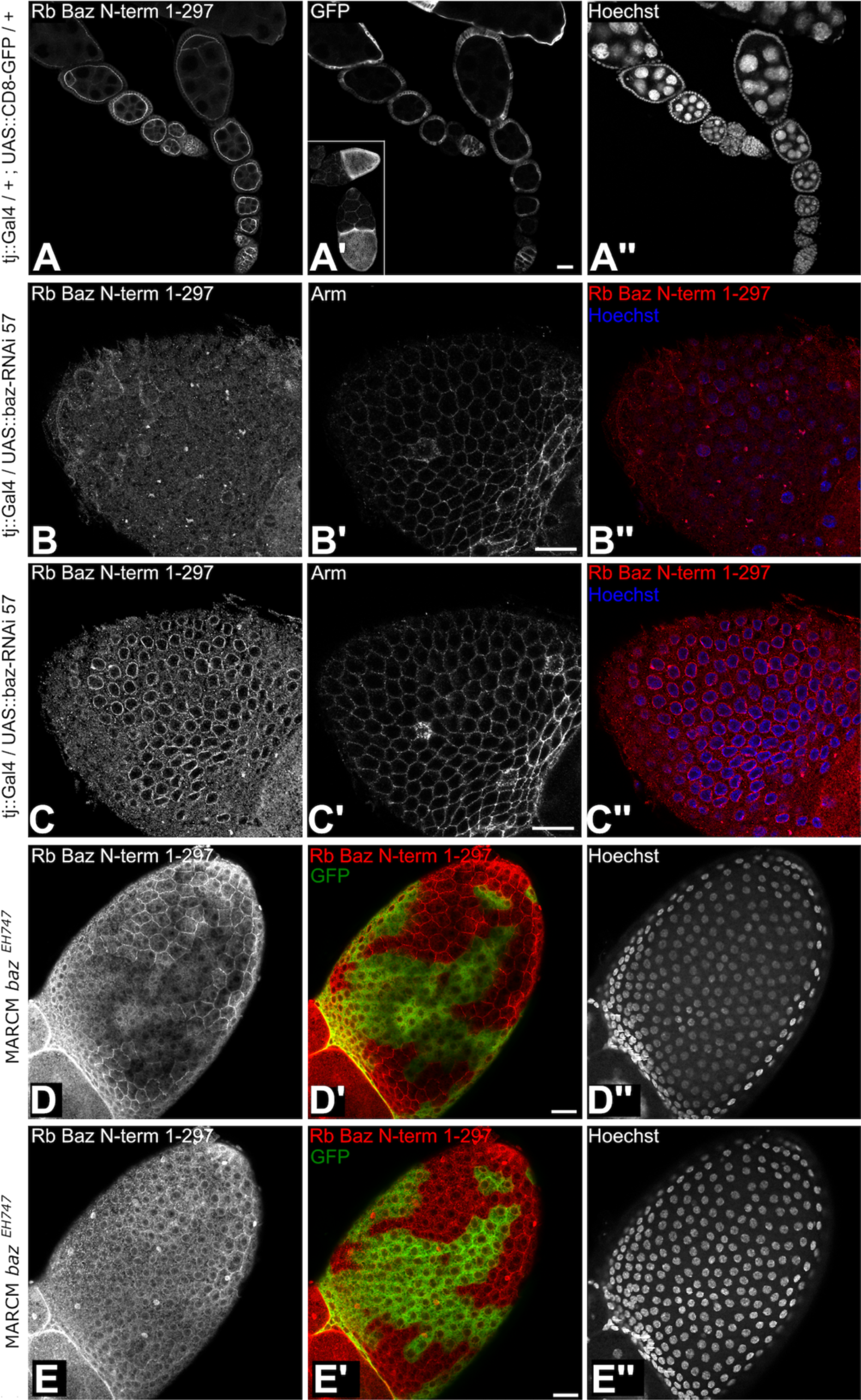
Nuclear envelope localization of Baz in follicular epithelial cells is a staining artifact. (A-A’’) Expression pattern of the tj::Gal4 driver line in follicular epithelial cells. Ovarioles of the genotype tj::Gal4/+; UAS::CD8-GFP/+ were stained with rabbit anti-Bazooka 1-297 (A), GFP (A’) and Hoechst (A’’). Anti-Baz immunostaining in early egg chambers shows an apical junctional signal in follicular epithelial cells (A). UAS::CD8-GFP under control of the tj-Gal4 driver shows GFP in all somatic follicular epithelial cells from early stages on (A’). At later stages GFP is expressed homogenously in follicle cells covering the oocyte (inset in A’). (B-C’’) baz-RNAi driven by tj::Gal4 leads to strong reduction of anti-Baz immunostaining at adherens junctions in the follicular epithelium (B) whereas junctional β-catenin (Arm) staining is clearly visible (B’). The merged image of anti-Baz immunostaining (red) and Hoechst DNA stain (blue) is shown in (B’’). (C-C’’) shows the same egg chamber as in (B-B’’) imaged at the level of the nuclei. Anti-Baz immunostaining is still detectable at the nuclear envelope (C) although there is no staining detectable at the junctions (compare to [B] and to Arm staining shown in [B’, C’]). The merged image of anti-Baz immunostaining (red) and Hoechst DNA stain (blue) is shown in (C’’). (D-E’’) MARCM loss-of-function clones for the null allele *baz^EH747^* in the follicular epithelium imaged at two different focal planes. (D-D’’) At the focal plane showing the adherens junctions, *baz^EH747^* homozygous mutant cells marked by GFP (green, D’) show a complete loss of junctional immunostaining with the anti-Baz antibody (D, D’). (E-E’’) In the focal plane of the nuclei, anti-Baz immunostaining shows prominent nuclear envelope staining in *baz^EH747^* homozygous mutant cells marked by GFP (green, E’) although these cells have lost junctional staining for Baz. (D’’, E’’) Hoechst staining for nuclei to show different focal planes. Scale bars = 20 µm.

For clonal analysis the null allele *baz^EH747^* was used, where a point mutation results in a premature stop after 51 amino acids. In follicle cells loss of Baz has no relevance to the junctions and further development of the egg chamber (Shahab et al., 2015). FLP-FRT and MARCM clones were generated in follicle cells and in the germline. Both small and large clones in the follicle epithelium showed the loss of Baz staining at the junctions (Fig. 2D, D’) but a persistent immunostaining signal for Baz around the nucleus (Fig. 2E, E’). Large follicle cell clones encompassing all follicle cells of an egg chamber marked by the loss of nuclear GFP (Fig. S4B’) showed the loss of junctional Baz in the *baz^EH747^* mutant follicle cells (Fig. S4B’’). The loss of Baz in the follicle cells did not result in any apparent junctional defect and the junctional marker Arm was still localized as in wild type (Fig. S4B’’’). In the germ line Baz was detectable at the junctions between the germ line cells, in particular between nurse cells and the oocyte. In addition, a strong signal was consistently detected at the nuclear envelope of the oocyte (Fig. S4A’’). While in *baz^EH747^* mutant germ line clones marked by the loss of nuclear GFP (Fig. S4C’), the junctional staining within the germline was lost, the staining around the oocyte nucleus and the nurse cell nuclei persisted (Fig. S4C’’). The staining for Arm revealed that the junctions between the germline cells were still intact (Fig. S4C’’’). The staining around the oocyte nucleus and the nurse cell nuclei was not detectable with the anti GFP antibody in the Baz-GFP BAC line (Fig. S4D’) nor in the Baz-GFP protein-trap line (Fig. S4E’). Staining with anti Baz again showed the oocyte nuclear envelope marked in these egg chambers (Fig. S4D’’, E’’).

### Baz nuclear envelope and NMJ immunostaining in the L3 body wall muscle does not reflect the true Baz localization

In parallel to our observations in the ovary we looked for Baz nuclear localization in the larval body wall muscle. Baz immunostaining was detectable at the nuclear envelope and at the neuromuscular junction (NMJ) as published before (Ruiz-Canada et al., 2004; Speese et al., 2012) (Fig. 3A - A’’). To test if this signal was specific for the antibody, we stained with the pre-immune serum, revealing a striped pattern in the muscle, but no signal at the NMJ or the nucleus (Fig. 3B - B’’). Staining with the secondary antibody only without any primary antibody resulted in no signal at the NMJ or the nucleus (Fig. 3C - C’’). Anti Discs large antibody and Hoechst were used in the same stainings as control to detect the NMJs and the nuclei (Fig. 3A - C’’). Together, these experiments showed that the Baz antibody was indeed responsible for the signal at the NMJ and the nuclear envelope of somatic body wall muscles.

**Figure 3.**
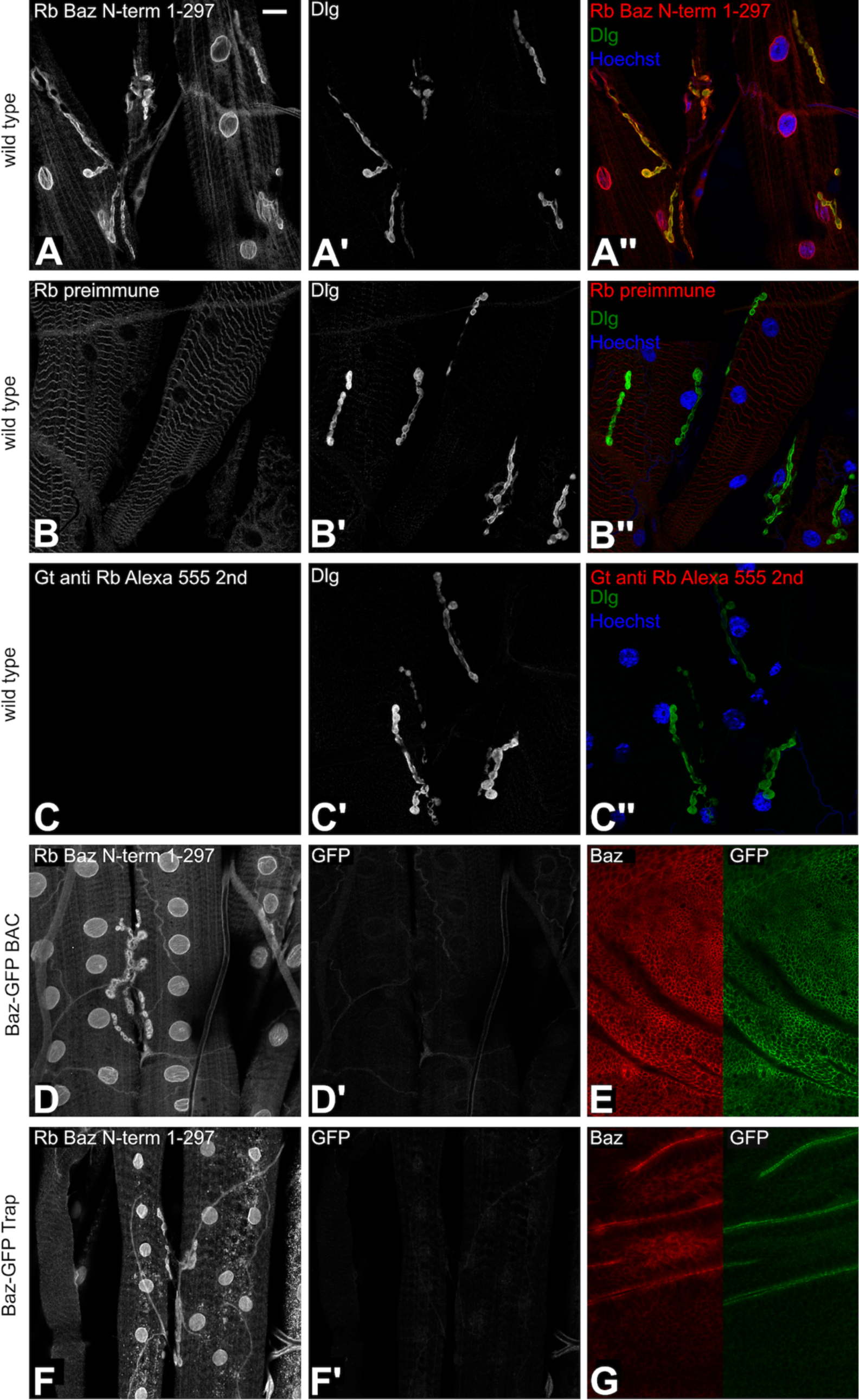
In larval body wall muscles staining of the nuclear envelope and the NMJ using the anti-Baz antibody is not reflected by the subcellular localization of Baz-GFP in the Baz-GFP BAC and gene trap lines. (A-A’’) Larval body wall muscles stained with anti-Baz antibody (A) and Dlg antibody (A’) to mark the neuromuscular junctions (NMJ). Anti-Baz immunostaining marks the nuclear envelope and the NMJs (A). The merged image is shown in (A’’). (B-B’’) Antibody staining with the pre-immune serum of the rabbit in which the anti-Baz antibody had been raised (B). Neither the nuclear membrane nor the NMJs are stained (B). Dlg staining of the NMJ is unaffected (B’). The merged image is shown in (B’’). (C-C’’) As a negative control, staining without primary rabbit antibody was performed, resulting in no signal (C). Dlg staining of the NMJ is unaffected (C’). The merged image is shown in (C’’). (D-E) Antibody staining of body wall muscle (D, D’) and wing imaginal disc (E) of a larva carrying the Baz-GFP BAC construct. (D) Anti- Baz immunostaining shows a signal at the nuclear envelope and the NMJ. (D’) GFP staining does not mark any structure in the body wall muscle. (E) Staining of a wing imaginal disc of the same larva as in (D) shows colocalization of Baz and GFP signals at epithelial junctions. (F-G) Antibody staining of body wall muscle (F, F’) and wing imaginal disc (G) of a larva of the Baz-GFP protein- trap line. (F) Anti-Baz immunostaining shows a signal at the nuclear envelope and the NMJ. (F’) GFP staining does not mark any structure in the body wall muscle. (G) Staining of a wing imaginal disc of the same larva as in (F) shows colocalization of Baz and GFP signals at epithelial junctions. Scale bar in (A) = 20 µm, valid for all panels.

By contrast, we did not detect GFP at the NMJ or at the nuclear envelope in larval body wall muscles (Fig. 3D’, F’) in the Baz-GFP BAC and the Baz-GFP-trap lines, whereas we observed a strong signal with the anti Baz antibody in these lines (Fig. 3D, F). In the wing imaginal discs of the same L3 larvae we observed colocalization of Baz immunostaining and Baz-GFP at intercellular junctions as expected (Fig. 3E, G).

It has been published that heterozygous *baz^4^* mutant larvae show a strong decrease in immunofluorescence signal of Baz and also of Spectrin at the NMJ (Ruiz-Canada et al., 2004). Another publication used the *baz^815-8^* allele to show a strong decrease in Baz and Spectrin at the NMJ (Ramachandran et al., 2009). However, both these *baz* alleles are likely to carry a second site hit as it has been shown that these alleles show a much stronger phenotype than a clean *baz* null allele in the follicular epithelial cells of the ovary (Shahab et al., 2015).

To reassess a potential localization of Baz at NMJs, we downregulated Baz by RNAi. Clonal analysis using a null allele of *baz* is not feasible in the muscle due to the syncytial nature of this tissue. To test whether RNAi works in body wall muscles we conducted *β-spectrin*-RNAi using the muscle specific (M12) driver line 5053-Gal4. We observed a strong decrease of α-spectrin staining in the postsynaptic membrane within the M12 muscle as published before (Pielage et al., 2006) (Fig. 4A - A’’). We pursued the same approach to downregulate Baz in muscle M12. The functionality of the *baz*-RNAi lines was demonstrated in the follicular epithelium, where expression under the control of tj::Gal4 led to strong downregulation of Baz at epithelial junctions (Fig. 2B). While it has been proposed that Baz is localized at the postsynaptic membrane within the NMJ (Ruiz-Canada et al., 2004), we did not detect any decrease in immunoreactivity for Baz or α-Spectrin upon *baz*-RNAi in M12 compared to other muscle segments not expressing *baz*- RNAi (Fig. 4B-B’’, C-C’’, compare NMJs marked by arrowhead and arrow). Baz nuclear envelope localization remained unaffected as well. We also did not see any downregulation of Baz or α- spectrin upon *baz*-RNAi in M12 at 29°C, when the UAS-Gal4 system is maximally active (data not shown). Taken together, our data strongly indicate that the nuclear envelope and NMJ immunostaining pattern observed using anti-Baz antibodies does not reflect the true Baz localization.

**Figure 4.**
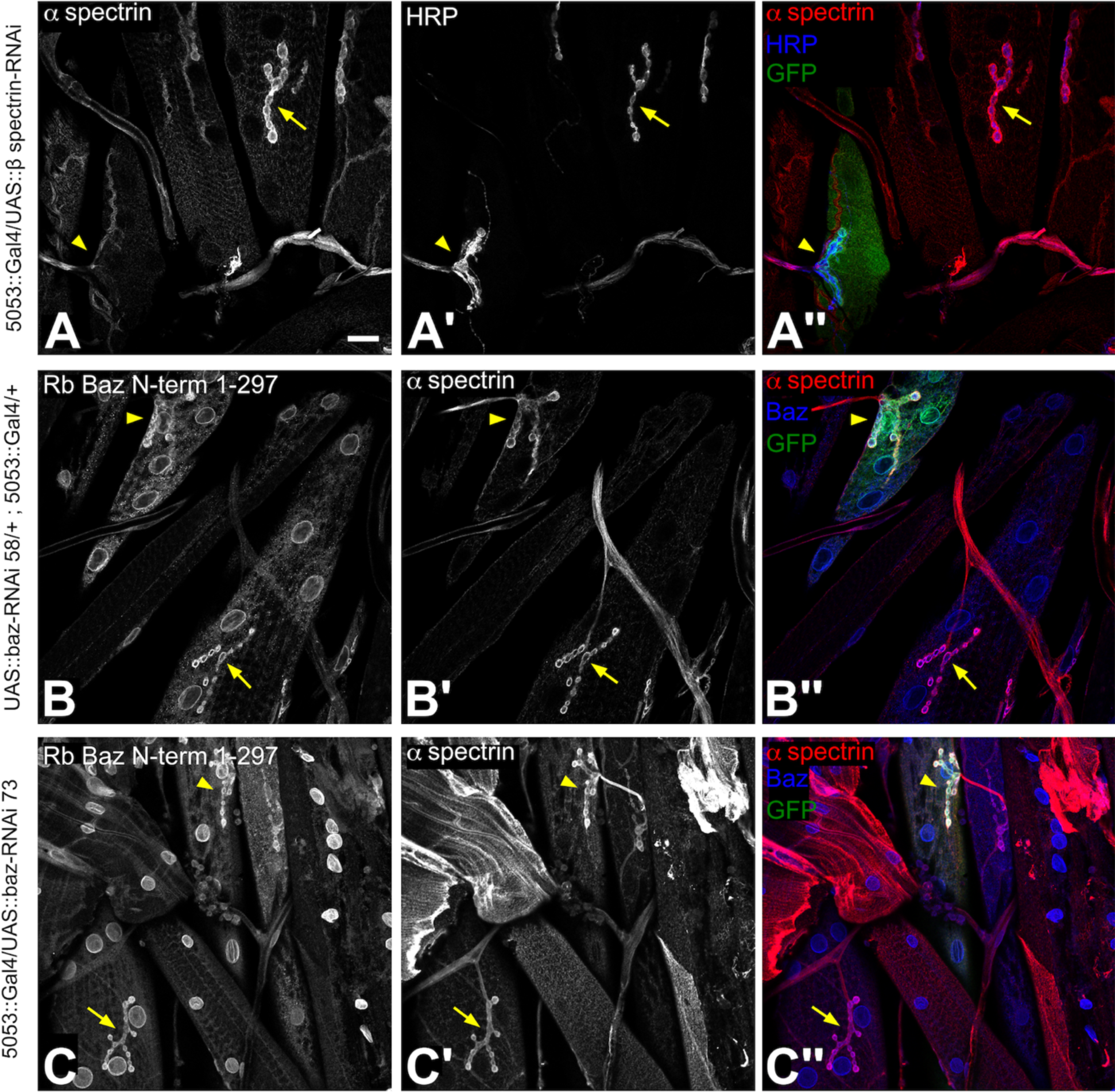
Nuclear envelope and NMJ localization of Baz in larval body wall muscles is a staining artifact. (A-A’’) β-Spectrin-RNAi was driven in muscle M12 by the 5053::Gal4 line. Larval body wall muscles were stained with antibodies against α-spectrin (A), HRP (A’) and GFP (green in merged image [A’’]) Co-expression of GFP marks muscle M12 (arrowhead), where the β- Spectrin-RNAi construct is driven (A’’). The arrow points to a control muscle not expressing β- Spectrin-RNAi to compare the NMJs and nuclei. (A, A’’) RNAi against β-Spectrin results in downregulation of α-Spectrin in the NMJ of muscle M12 (arrowhead). In the neighboring control muscle α-Spectrin is expressed at synaptic boutons (arrow). (A’) HRP staining to mark the NMJs. (B-C’’) Baz-RNAi with two different constructs (baz-RNAi 58 and baz-RNAi 73) does not show downregulation of anti-Baz immunostaining at the nuclear envelope or the NMJ in M12 (arrowhead) in comparison to a control muscle not expressing baz-RNAi (arrow). (B’, C’) α- Spectrin is also not downregulated in the NMJ of muscle M12 (arrowhead) in comparison to control (arrow). Merged images with GFP (green) are shown in (B’’, C’’). Scale bar in (A) = 20 µm, valid for all panels.

## Discussion

### Baz nuclear envelope localization is an artifact of immunostaining

In line with results from (Speese et al., 2012), our initial immunostaining analysis revealed strong localization of Baz to the nuclear envelope and colocalization with the nuclear envelope protein Lamin C. The fact that four different anti Baz antibodies raised against three different regions of the Baz protein showed nuclear localization provided strong evidence to believe that the observed immunostaining pattern was unlikely to be an immunostaining artifact. Further analysis revealed that localization of Baz to the nuclear envelope was detectable only in specific cell types. We found Baz nuclear envelope immunostaining mostly in polyploid cells undergoing endoreduplication such as larval fat body cells, larval body wall muscle, nurse cells and late-stage follicular epithelial cells, but also in diploid oocytes and spermatogonia.

However, analysis of Baz immunostaining in mutant clones for two different *baz* null alleles indicated that the observed nuclear envelope localization pattern was not specific to Baz. We initially speculated whether the persistence of Baz nuclear envelope immunostaining in the *baz* mutant follicular epithelial cells and oocyte could be due to increased Baz protein stability and low turnover of Baz associated with the nuclear envelope. However, this is unlikely given that no change in the intensity of Baz nuclear envelope staining was observed when large late-stage follicular epithelial clones were compared to small early-stage clones. If perdurance would be the explanation for the persistent nuclear envelope staining, then the signal should become weaker as cells divide and the tissue expands. Similarly, in *baz* mutant nurse cells, Baz staining at the junctions was completely lost whereas no change in nuclear envelope staining intensity was observed.

No GFP tagged Baz protein from the Baz-GFP-trap line or the Baz-GFP BAC line was detected at the nuclear envelope by immunofluorescence analysis with or without GFP antibody immunostaining. Thus, our data strongly indicate that Baz does not localize to the nuclear envelope and that nuclear envelope immunostaining observed with anti-Baz antibodies is due to cross reactivity with an unidentified epitope. We can only speculate about the reason for the nuclear envelope signal detected with the different anti Baz antisera. All four sera were raised against GST fusion proteins, so it could be that antibodies against GST that are present in the antisera cross-react with a component of the nuclear envelope. However, as we were interested in a potential function of Baz at the nuclear envelope, we did not further investigate this after we found that signal to be unrelated to Baz.

### Baz does not localize to the postsynaptic membrane of the neuromuscular junction

Finally, our data also revealed that the published localization of Baz to the postsynaptic membrane of the NMJ (Ramachandran et al., 2009; Ruiz-Canada et al., 2004; Speese et al., 2012) is an artifact, as the staining is unaffected by RNAi against Baz and neither the GFP-Baz trap line nor the GFP-Baz BAC line showed any GFP signal at the NMJ. Surprisingly though, work from the Budnik lab reported NMJ phenotypes such as reduced staining intensity for α-Spectrin and reduced number of synaptic boutons in animals heterozygous mutant for the *baz* alleles *baz^4^* and *baz^815-8^* as well as upon RNAi against *baz* (Ramachandran et al., 2009; Ruiz-Canada et al., 2004). We did not attempt to reproduce these findings because our data do not provide evidence for localization of Baz at the NMJ. The reported effects on the NMJ may be explained by the existence of second site mutations on the chromosomes carrying the *baz^4^* and *baz^815-8^* alleles used for the analyses. These second site mutations apparently also cause defects in epithelial apical-basal polarity that are not observed using clean null alleles of *baz* (Shahab et al., 2015).

Altogether, the reported findings for a function of Baz at the nuclear envelope (Speese et al., 2012) and at the NMJ (Ramachandran et al., 2009; Ruiz-Canada et al., 2004) should be regarded with great caution as we did not find any evidence for localization of Baz to these subcellular structures.

## Materials and Methods

### Fly stocks and genetics

The following fly stocks were used in this study: *w^1118^* (was used as wild type control, BL 3605), Baz-GFP BAC (Besson et al., 2015), Baz-GFP gene trap CC01941 (BL 51572) (Buszczak et al., 2007) UAS::Baz-RNAi57 (VDRC v2915), UAS::Baz-RNAi58 (VDRC v2914), UAS::Baz-RNAi73 HMS01412 (BL 35002), UAS::β-spectrin-RNAi (Pielage et al., 2005), UAS::CD8-GFP (BL 32184), CY2::Gal4 (Queenan et al., 1997), Traffic Jam::Gal4 (Li et al., 2003), 5053::Gal4 M12 (BL 2702), *FRT19A* (BL 1709), *baz^EH747^* FRT19A, *baz^4^* FRT19A, *baz^XR11^* FRT19A, (Shahab et al., 2015), *hsFlp^122^ FRT19A H2AvD-GFP* (BL 32045), *FRT19A tubP::Gal80LL1 hsFLP; tubP::Gal4 UAS::mCD8-GFP* (for generation of MARCM clones, gift from Heinrich Reichert). Stock numbers from the Bloomington *Drosophila* Stock Center (BL #) and from the Vienna *Drosophila* Research Center (VDRC #) are given in parentheses. Crossings for RNAi experiments were set up at 25°C or 29°C. For generating clones in ovaries by Flipase-mediated mitotic recombination of the FRT sites flies were heat shocked for 1h at 37°C 5-7 days prior to preparation of the ovaries.

### Preparation of Ovaries

One day before preparation the females were fed with fresh yeast. Ovaries were removed and ovariole tubules separated by pipetting the ovaries prior to fixation two times with a cut 1000µl tip to enlarge tip opening. Ovaries were fixed in 3,7% formaldehyde, washed three times in PBS and blocked in PTX (0,1% Trixon X-100 in PBS) for at least 3h.

### Preparation of larval muscle

L3 larvae were washed in PBS to remove residual food and placed on a plate with PBS. Larvae were fixed with needles and cut open between the lateral trunks of the tracheae along the a/p- centerline. Larval cuticle was nicked at the anterior and posterior end to the lateral side and the cuticle was unfolded to fix it with needles onto the plate to flatten it. Inner organs were removed, imaginal discs remained. Torsos were washed with PBS to remove residual tissue and then shortly once with 3,7% formaldehyde in PBS and then fixed for 15 min in 3,7% FA. Fixative was removed and the torso washed three times with PBS. After the needles were removed, the torsos were transferred to a tube and washed three times with PTX (0,1% Triton X-100 in PBS) for 10min.

### Immunostainings and antibodies

The following antibodies were used for immunostainings: rabbit anti Baz-N-term (aa 1-297) (Wodarz et al., 2000) 1:1000, rabbit anti Baz-N-term (aa1-297) pre-immune 1:1000, rat anti Baz- N-term (aa 1-297) (Wodarz et al., 1999) guinea pig anti Baz-PDZ (aa 309-747) (Shahab et al., 2015) 1:1000, guinea pig anti Baz-C-term (aa 905-1221) 1:1000 (this work), mouse anti α- Spectrin (3A9, DSHB) 1:10, mouse anti Dlg (4F3, DSHB) 1:20, mouse anti Armadillo (N2-7A1, DSHB) 1:20, mouse anti Lamin C (ADL 76.10, DSHB) 1:100, rat anti DE-Cadherin (DCAD2, DSHB) 1:5, goat anti HRP-Alexa647 (Jackson ImmunoResearch), mouse anti GFP (A11120 Molecular Probes) 1:1000, rabbit anti GFP (A11122 Molecular Probes) 1:1000. Secondary antibodies conjugated to Alexa-Fluor-488/555/647 (Invitrogen) were used at 1:400. DNA was stained with HOECHST 33258 (Sigma). Immunostainings for the 1st and 2nd antibodies were performed in 5% NHS in PTX. Stained tissues were imaged on a Zeiss LSM880 Airyscan confocal microscope.

## Acknowledgements

We thank Uli Thomas for showing us how to prepare and stain larval muscles, Jan Pielage, Trudi Schüpbach, Dorothea Godt, Francois Schweisguth, Heinrich Reichert, the Bloomington *Drosophila* stock center at the University of Indiana, the Vienna *Drosophila* Research Center (VDRC) and the Developmental Studies Hybridoma Bank (DSHB) at the University of Iowa for sending fly stocks and reagents. We thank Mona Honemann-Capito, Ferdi Grawe and Monique Ulepic for technical assistance and members of the Wodarz lab for discussion. This work was funded by grants from the Deutsche Forschungsgemeinschaft (Cluster of Excellence 171 “Nanoscale Microscopy and Molecular Physiology of the Brain”).

---

**Figure S1.**
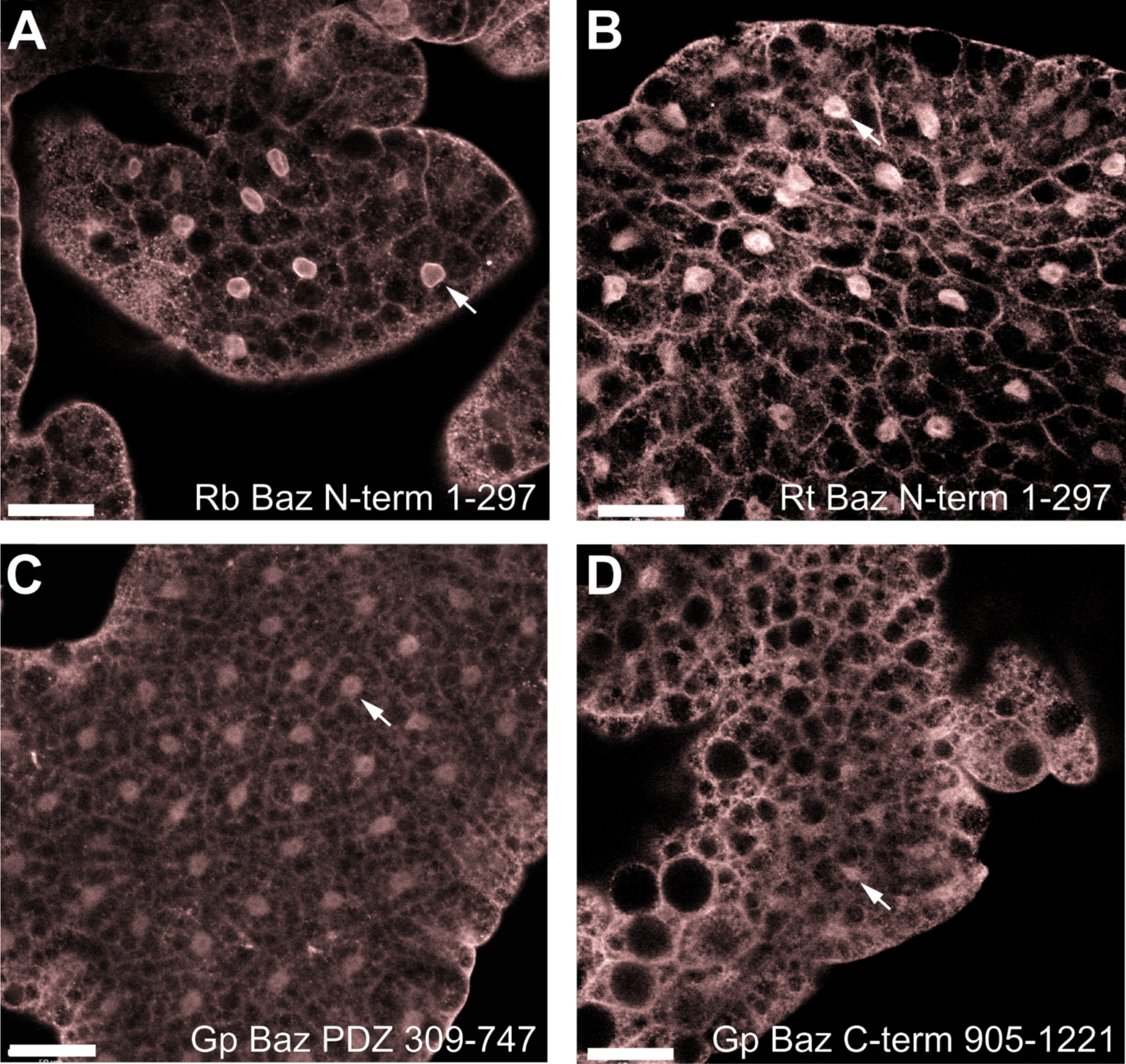
Baz nuclear envelope localization is observed with multiple antibodies raised against non-overlapping domains of Baz. (A-D) Fat body tissue dissected from wild type 3^rd^ instar larvae and stained with different anti-Baz antibodies. Rabbit anti-Baz N-term 1-297 (A) and Rat anti-Baz N-term 1-297 (B) were raised against a GST fusion protein containing amino acids 1-297 from the N-terminal region of Baz. Guinea pig anti-Baz PDZ 291-737 (C) was raised against a GST fusion protein containing amino acids 291-737 of Baz corresponding to the PDZ domains. Guinea pig anti-Baz C-term 905-1221 (D) was raised against a GST fusion protein containing amino acids 905-1221 of Baz. Arrows indicate nuclei. Scale bars = 50 µm.

**Figure S2.**
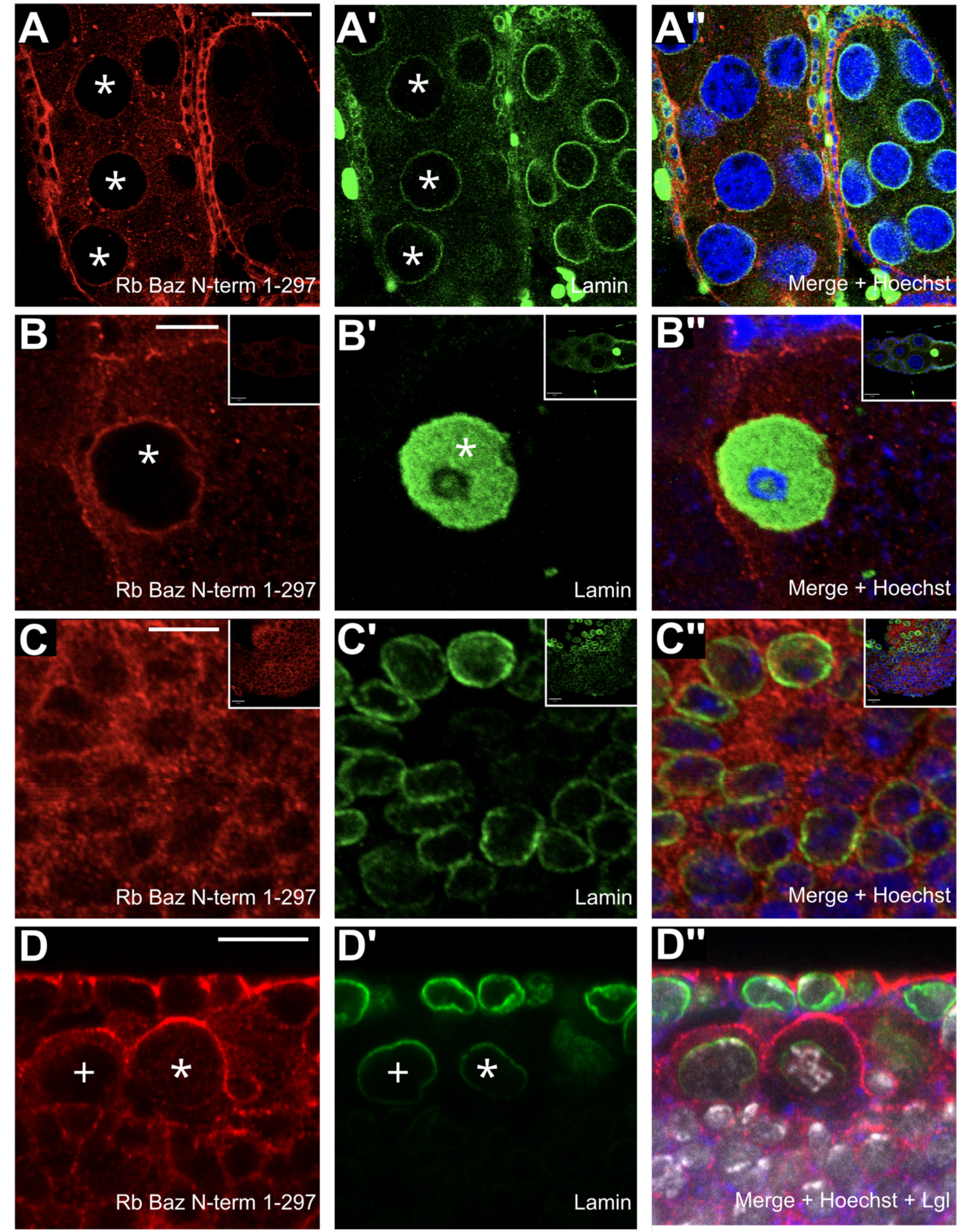
Anti-Baz immunostaining colocalizes with Lamin C at the nuclear envelope of nurse cells and the oocyte but not in larval imaginal disc cells, embryonic epidermal cells and neuroblasts. (A-A’’) Adult ovaries were stained with anti-Baz (A), Lamin C (A’) and Hoechst (blue in merged image [A’’]). The images show two egg chambers with the focus at the plane of the large nurse cell nuclei (asterisks). (B-B’’) High magnification images of the oocyte nucleus (asterisk) in a stage 8 egg chamber stained with anti-Baz (B), Lamin C (B’) and Hoechst (blue in merged image [B’’]). The inset shows a low magnification image of the whole egg chamber. (C-C’’) Detail of a third instar wing imaginal disc stained with anti-Baz (C), Lamin C (C’) and Hoechst (blue in merged image [C’’]). The images were taken at the focal plane of the nuclei. No enrichment of anti-Baz immunostaining is visible at the nuclear envelope. The inset shows a low magnification of the wing imaginal disc. (D-D’’) Sagittal section of a stage 8 embryo stained with anti-Baz (D), Lamin C (D’), Hoechst (gray scale in merged image [D’’]) and Lgl (blue in merged image [D’’]). An interphase neuroblast is marked by (+) and a prophase neuroblast is marked by (*). Lgl serves to mark the basolateral cortex of the epidermal epithelium (top). Anti-Baz immunostaining marks the apical adherens junctions of the epidermis and the cortex of neuroblasts, with enrichment in the apical cortex in the prophase neuroblast, but is absent from the nuclear envelope (D). Scale bar in (A) = 20 µm, Scale bars in (B) and (C) = 5 µm, Scale bar in (D) = 10 µm.

**Figure S3.**
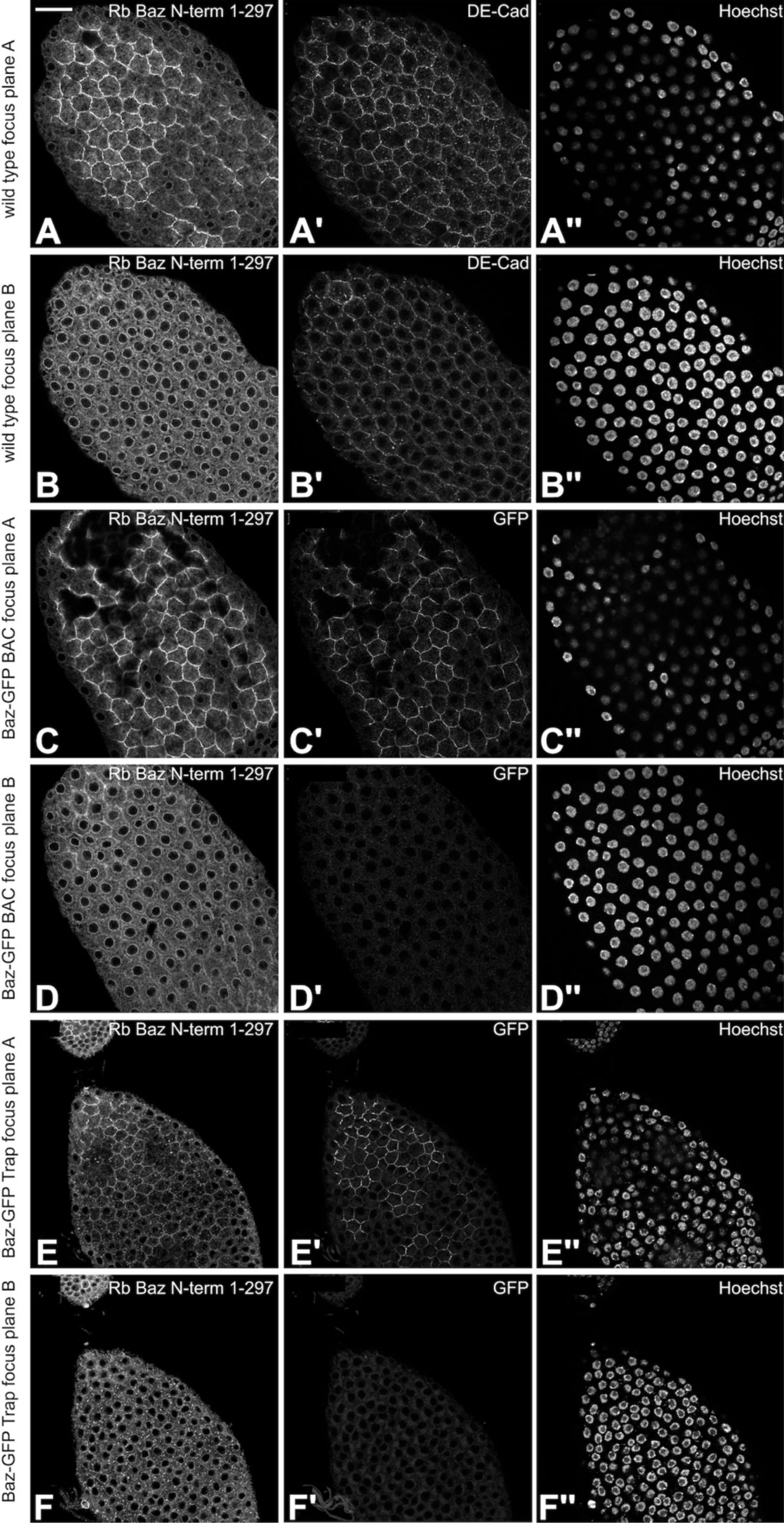
Anti-Baz immunostaining colocalizes with Baz-GFP in epithelial junctions of the follicular epithelium but not at the nuclear envelope. (A-B’’) Follicular epithelium of a wild type egg chamber imaged at the focal plane of the adherens junctions (A-A’’) and the nuclei (B-B’’) stained with anti-Baz (A, B), DE-Cad (A’, B’) and Hoechst (A’’, B’’). Junctional staining of Baz colocalizes with DE-Cad (A, A’) whereas the nuclear envelope staining of Baz does not show any overlap with DE-Cad staining (B, B’). Hoechst staining in (A’’, B’’) is shown to demonstrate the focal plane of the respective images. (C-D’’) Follicular epithelium of the Baz-GFP BAC line imaged at the focal plane of the adherens junctions (C-C’’) and the nuclei (D-D’’) stained with anti-Baz (C, D), GFP (C’, D’) and Hoechst (C’’, D’’). Junctional staining of anti-Baz colocalizes with Baz-GFP (C, C’) whereas the nuclear envelope signal of anti-Baz does not show any overlap with Baz-GFP staining, which is barely detectable at the focal plane of the nuclei (D, D’). Hoechst staining in (C’’, D’’) is shown to demonstrate the focal plane of the respective images. (E-F’’) Follicular epithelium of the Baz-GFP protein-trap line imaged at the focal plane of the adherens junctions (E-E’’) and the nuclei (F-F’’) stained with anti-Baz (E, F), GFP (E’, F’) and Hoechst (E’’, F’’). Junctional staining of anti-Baz colocalizes with Baz-GFP (E, E’) whereas the nuclear envelope signal of anti-Baz does not show any overlap with Baz-GFP staining, which is barely detectable at the focal plane of the nuclei (F, F’). Hoechst staining in (E’’, F’’) is shown to demonstrate the focal plane of the respective images. Scale bar in (A) = 20 µm, valid for all panels.

**Figure S4.**
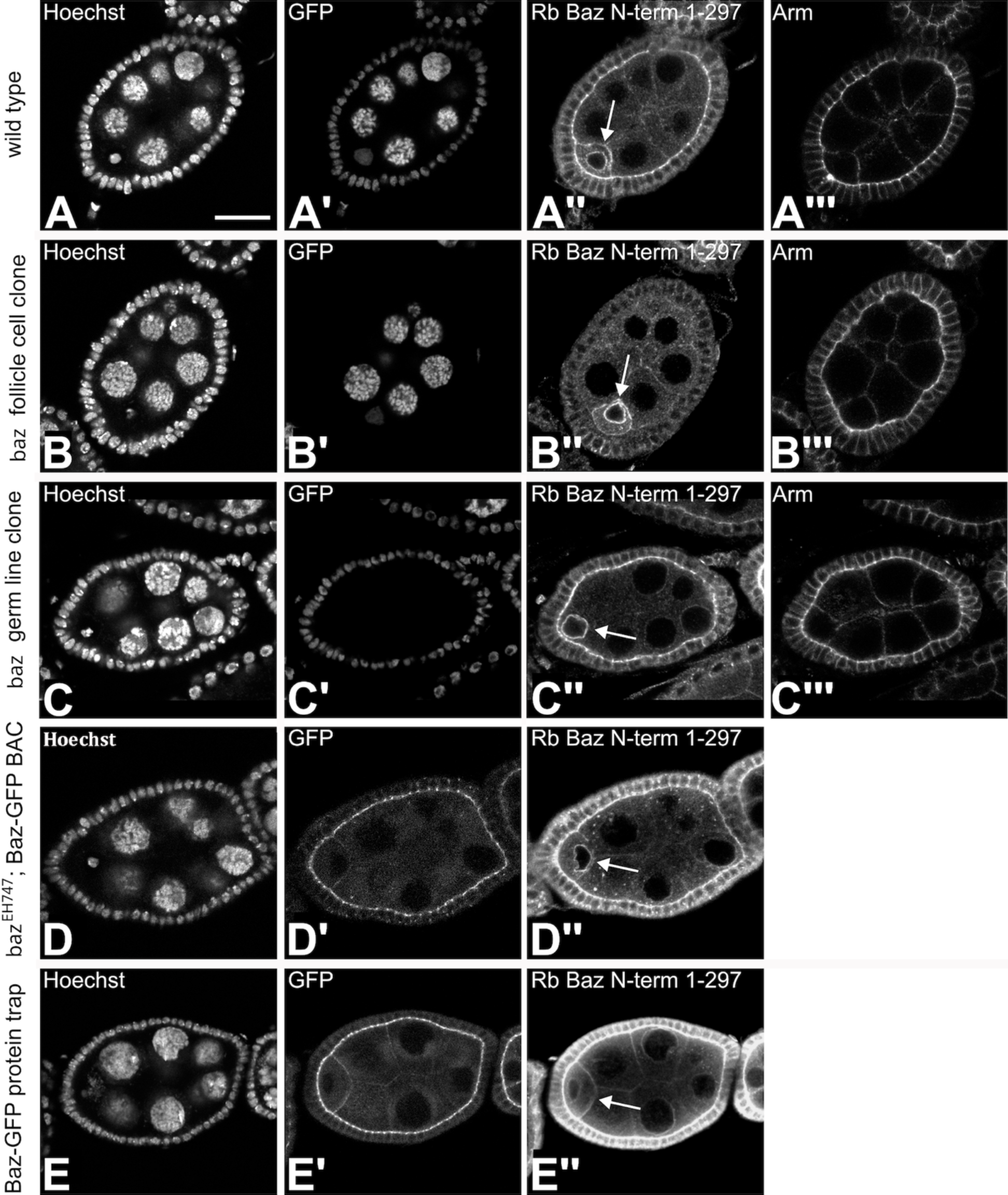
Nuclear envelope localization of Baz at the oocyte nucleus is a staining artifact. (A-A’’’) Wild type egg chamber expressing nuclear GFP at stage 4/5 with the focal plane at the oocyte nucleus stained with Hoechst (A), GFP (A’), anti-Baz (A’’) and Arm (A’’’). Anti-Baz immunostaining marks the apical adherens junctions of the follicular epithelium, the junctions between the germline cells and the nuclear membrane of the oocyte (A’’, arrow). Immunostaining of Arm marks the apical junctions and the lateral membrane of the follicular epithelial cells and junctions between germline cells (A’’’). (B-B’’’) Large follicle cell clone of *baz^EH747^*. All nuclei are marked by Hoechst staining (B). Loss of nuclear GFP shows that the complete follicular epithelium is homozygous mutant for *baz^EH747^* (B’). Note that anti-Baz immunostaining is lost at the apical junctions of follicular epithelial cells, but still visible in the germline, especially at junctions between oocyte and nurse cells and the nuclear envelope of the oocyte (B’’, arrow). Junctional staining for Arm is unaffected by loss of Baz in follicle cells (B’’’). (C-C’’’) Germ line clone of *baz^EH747^*. All nuclei are marked by Hoechst staining (C). Loss of GFP in germ line nuclei marks the *baz^EH747^* mutant germ line clone (C’). Anti-Baz immunostaining marks the apical adherens junctions of the follicular epithelium, whereas the signal at the junctions between the germline cells is lost. However, the signal at the nuclear envelope of the oocyte and the nurse cells persists (C’’, arrow). Junctional staining for Arm is unaffected by loss of Baz in germ line cells (C’’’). (D-D’’) Comparison of the signal using anti-Baz and anti-GFP antibodies in the Baz-GFP BAC line in the *baz^EH747^* mutant background. While anti-Baz immunostaining shows a signal at the nuclear envelope of the oocyte (arrow) and the nurse cells in addition to junctional staining in the follicular epithelium and in between germ line cells (D’’) the GFP antibody does not detect a signal at the nuclear envelope of the oocyte but only at the junctions in the follicular epithelium and in between the germ line cells (D’). (E-E’’) Immunostaining using anti-Baz and anti-GFP in the Baz-GFP protein-trap line. Results are identical to those shown for the Baz-GFP BAC line. Scale bar in (A) = 20 µm, valid for all panels.

**Table 1.** Assessment of nuclear staining patterns for different Baz antibodies.

| **Anti-Baz Antibody** | **Nuclear staining pattern** |
| --- | --- |
| Rabbit anti-Baz DE99646 (a.a 1-297 N-term) | Nuclear envelope localization |
| Rat anti-Baz DE99647-1 (a.a 1-297 N-term) | Nuclear envelope localization |
| G.P anti-Baz (a.a 905-1221 C-term) | Weak staining, some nuclear localization |
| G.P anti-Baz (a.a 291-737 PDZ domains) | Weak staining, some nuclear localization |

## Notes

### Competing Interest Statement

The authors have declared no competing interest.

